# Kinetic and structure-based comparisons of silent and stimulatory flagellin interactions with TLR5

**DOI:** 10.1101/2024.12.09.627598

**Authors:** Michael E. W. Bell, Miriam Haag, Sara J. Clasen, Marieke Böcker, Kateryna Maksymenko, Iris Koch, Katharina Hipp, Marcus D. Hartmann, John R. Weir, Felipe Merino, Ruth. E. Ley

**Affiliations:** Department of Microbiome Science, Max Planck Institute for Biology, Tübingen, Germany; Cluster of Excellence EXC 2124 Controlling Microbes to Fight Infections, University of Tübingen, Tübingen, Germany; Department of Protein Evolution, Max Planck Institute for Biology, Tübingen, Germany; Electron Microscopy Facility, Max Planck Institute for Biology, Tübingen, Germany; Friedrich Miescher Laboratory of the Max Planck Society, Tübingen, Germany; Interfaculty Institute of Biochemistry, University of Tübingen, Tübingen, Germany; German Center for Infection Research (DZIF), partner-site Tübingen, Tübingen, Germany

## Abstract

The bacterial protein flagellin is the sole ligand of the innate immune receptor Toll-like receptor 5 (TLR5). Flagellins with strong agonism bind TLR5 at their D1 and D0 domains, while poor agonist “silent” flagellins bind with D1 only. However, D0-TLR5 interactions insufficiently explain the silent phenotype. Here, we characterize the D1 domain binding kinetics of the silent flagellin *Rh*FlaB compared to the strong canonical agonist *St*FliC. Using Surface Plasmon Resonance, we show that the *Rh*FlaB-D1 binds more strongly than *St*FliC-D1, but forms shorter-lived complexes. Cryo-EM analysis of *Rh*FlaB showed its D1 primary interface (PI) contains two distinct hydrophobic pockets, which should facilitate rapid association, while its D1’s secondary interface (SI) is dominated by negatively charged residues, which likely impede residue-residue interactions. Comparisons with an existing *St*FliC-D1 structure suggest that strong agonism requires strong binding at both the PI and SI of the D1, and silent flagellins evade detection partly through weaker SI interactions. These findings offer a foundation for designing targeted interventions that can modulate TLR5 activity to influence immunity, inflammation, or even gut microbiome composition.

## Introduction

Many gut-resident bacteria possess motility facilitated by flagella, whip-like structures composed of flagellin monomers, which increase their fitness by enabling them to reach preferred niches. The structure of flagellin is highly conserved across motile bacterial species, generally forming an L-shape composed of two highly conserved domains (D0 and D1), which form the core of the flagellum, and hypervariable domains (D2 and D3), which form the outer surface (Andersen-Nissen et al. 2005; Beatson, Minamino, and Pallen 2006; Kreutzberger et al. 2022).

Bacterial flagellins are recognised by human Toll-Like Receptor 5 (TLR5), a membrane bound innate immune receptor, triggering a pro-inflammatory cytokine response (Kumar, Kawai, and Akira 2009; Burgueño and Abreu 2020; Hayashi et al. 2001). The flagellin motifs targeted by TLR5 are located in the D0 and D1 domains, and are integral for polymerisation when forming the flagellum (Fig. 1A) (Kim et al. 2018; Smith et al. 2003; Song et al. 2017; Forstnerič et al. 2017; Yoon et al. 2012).

**Figure 1.**
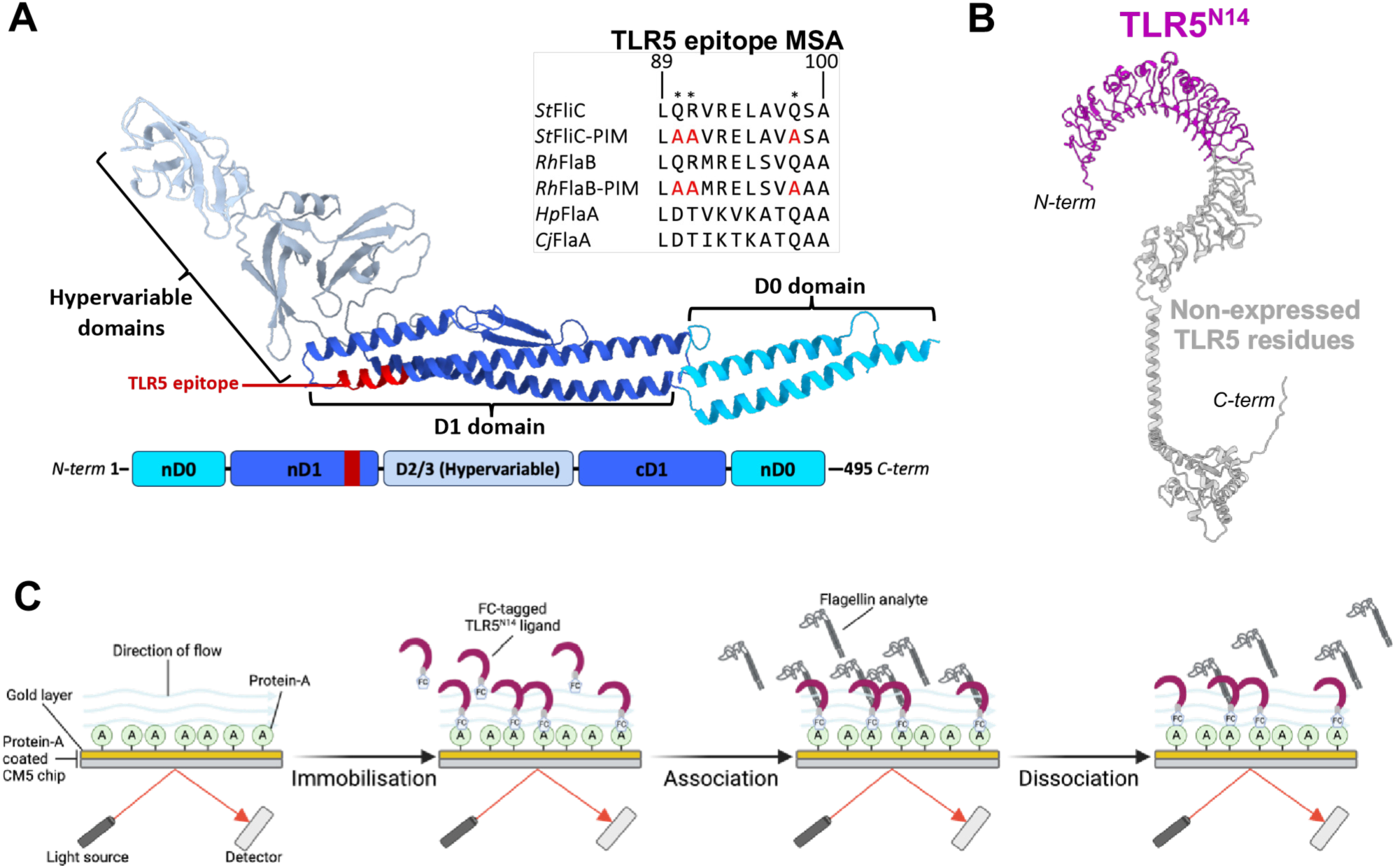
Overview of flagellin, TLR5^N14^, and the surface plasmon resonance methodology. (A) An *St*FliC structure showing the ‘standard’ flagellin domain layout (PDB ID #6jy0) and a ClustalO (1.2.4) multiple sequence alignment (MSA) of the TLR5 epitope in notable flagellins. The residues mutated in ‘PIM’ mutants are indicated with an ’*’. Each domain is coloured separately, corresponding to the domain map, with the TLR5-epitope highlighted in red. (B) The predicted structure of human TLR5 generated with Alphafold2 (Jumper et al. 2021). The region expressed in the TLR5^N14^ construct is coloured in purple, and residues not expressed are shown in grey. (C) Graphical overview of the SPR analysis methodology.

Previous studies on TLR5-flagellin interactions have focused primarily on flagellin produced by bacterial pathogens, such as FliC from *Salmonella enterica* subspecies. Yoon et al. identified a set of residues in the *Salmonella enterica* serovar *Dublin* FliC (*Sd*FliC) D1 domain important for TLR5 binding and necessary for signaling (Yoon et al. 2012). Evasive pathogens such as *Helicobacter pylori* and *Campylobacter jejuni* lack these key residues, fail to bind, and evade TLR5 recognition (Andersen-Nissen et al. 2005; Kim et al. 2018; Kreutzberger et al. 2020). Further mutagenic and structural studies have also identified a likely secondary interaction site in the D1 domain, though its role in TLR5-flagellin interactions has been poorly characterised (Song et al. 2017; Yoon et al. 2012; Smith et al. 2003; Ivičak-Kocjan et al. 2018).

Our recent work highlighted that many flagellins from commensal gut bacteria are poor TLR5 agonists, despite displaying a comparatively strong binding strength. We termed such flagellins “silent” (Clasen et al. 2023). In-depth comparative characterization of the stimulatory flagellin *St*FliC (produced by *Salmonella enterica* serovar Typhimurium) to the silent flagellin *Rh*FlaB (produced by the common commensal bacterium *Roseburia hominis*) revealed that *St*FliC has an additional allosteric binding site necessary for signalling. We mapped the allosteric binding site to the C-terminal region of the *St*FliC D0 domain. This allosteric binding site is lacking in the *Rh*FlaB D0 domain. Swapping in of the *St*FliC-D0 to *Rh*FlaB substantially rescued signalling, bringing them closer to *St*FliC levels. But the incomplete rescue of receptor activity suggested other uncharacterized aspects of the D0-domain binding may also tune the agonism of flagellins.

One way that the D0 may be involved in receptor response is through its binding kinetics. Temporal differences in complex formation could tune the TLR5 receptor response response by limiting the ability of the D0-domain to bind, and thereby reducing the timeframe for TLR5 signal propagation (Oda and Kitano 2006; Chen et al. 2001). However, although we had observed comparatively stronger binding in the *Rh*FlaB D1 domain to TLR5 compared the *St*FliC-D1 (Clasen et al. 2023), their binding kinetics remain to be elucidated and compared.

Here, we characterised the temporal nature of interactions between *St*FliC, *Rh*FlaB, and TLR5^N14^, a truncated human TLR5 construct containing only the D1-domain binding sites, using Surface Plasmon Resonance (SPR). Next, we determined the structure of *Rh*FlaB from sheared *R. hominis* flagellar filaments by electron cryo-microscopy (cryo-EM). Comparison to published *St*FliC structure data allowed us to identify structural features of the D1 domain which differed between the two flagellins. Taken together, results from these two approaches suggest a kinetics-based mechanism by which distinct surface characteristics of the silent flagellin D1 domain impairs complex stability, limiting the ability for TLR5 to trigger a downstream immune response.

## Results

### R. hominis FlaB displays stronger binding to TLR5^N14^ compared to S. typhimurium FliC

We used SPR to assess the binding kinetics of flagellins, including (i) wildtype StFliC, a canonical human TLR5 strong agonist produced by Salmonella enterica serovar Typhimurium, (ii) RhFlaB, a flagellin produced by the commensal human gut bacterium Roseburia hominis (Clasen et al. 2023), (iii) primary interface mutants (PIM) and (iv) ΔD0 mutants. PIM mutants were made with three substitutions in the TLR5 epitope (StFliC: Q89A, R90A, Q97A; RhFlaB: Q88A, R89A, Q96A)(Figure 1). These residues are directly involved in TLR5 D1 domain binding, and the substitutions cause drastically reduced binding and stimulatory activity (Smith et al. 2003; Kim et al. 2018). ΔD0 mutants are truncated to remove the D0 domain, which is not expected to interact with TLR5^N14^.

Our measures of binding strength (KD) indicate *Rh*FlaB binds TLR5 slightly but significantly more strongly than *St*FliC (KD 11.38 nM vs 16.26 nM) (Fig. 2A; Supp. Fig 1; Table 1), corroborating previous observations (Clasen et al. 2023). As expected from substitutions in the key binding residues, *Rh*FlaB-PIM (KD 77.99 nM) had a 7-fold increase in KD compared to *Rh*FlaB. Similarly, *St*FliC-PIM (KD 585.1 nM) showed a 36-fold increase in KD compared to *St*FliC. These greatly increased KD values indicate that the PIM substitutions severely impaired TLR5 binding in both flagellins (Table 1). Flagellin mutants lacking the D0 domain (ΔD0) showed marginally lower KD values compared to non-ΔD0 equivalents, but these differences were not statistically significant (Table 1; Supp. Fig. 1).

**Figure 2.**
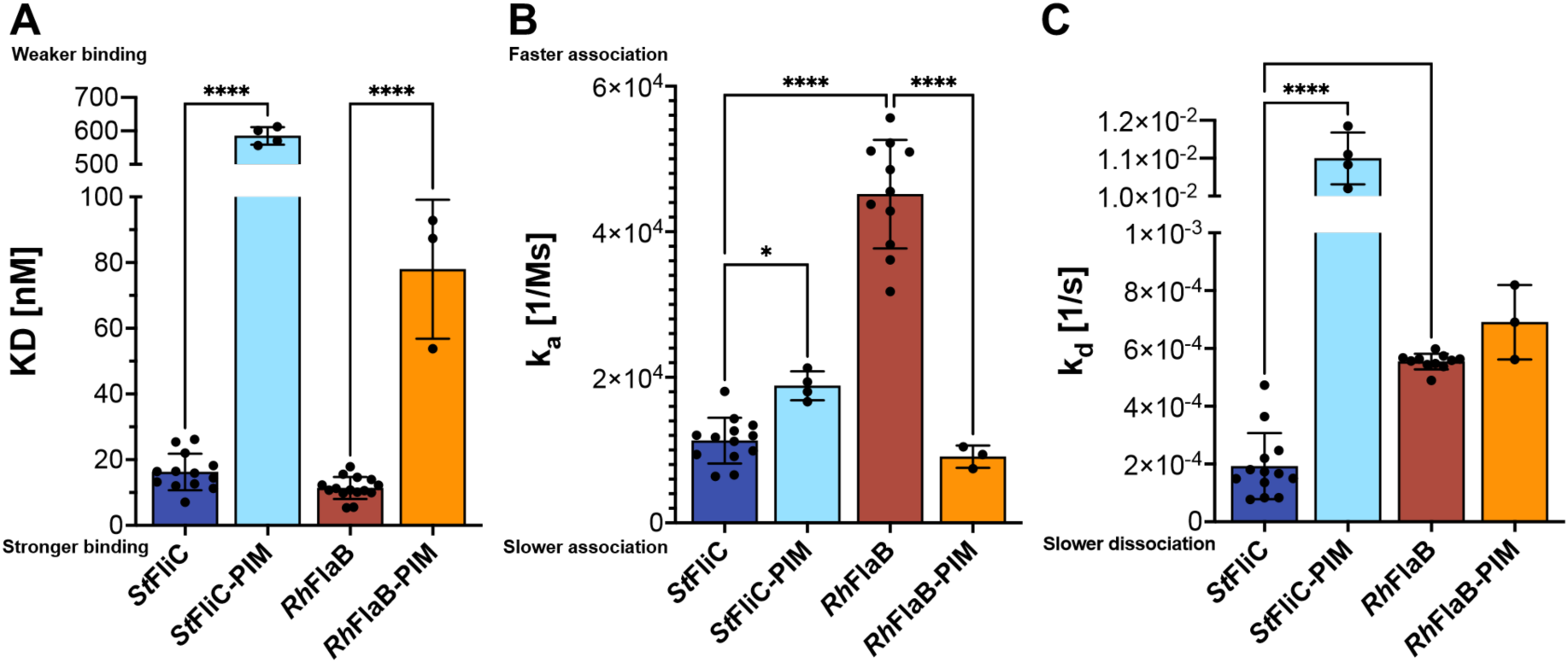
Differing interaction kinetics are observed in *St*FliC and *Rh*FlaB binding to TLR5. Pooled SPR binding (A) and kinetics (B-C) results comparing *St*FliC and *Rh*FlaB to their respective PIM mutants. Each bar includes data from at least 3 replicates. Error bars represent SD, significance was calculated by ordinary one-way ANOVA with Šídák correction in Prism 10 (GraphPad) (\**p* < 0.05, \*\**p* < 0.01, \*\*\*\**p* < 0.0001).

**Table 1.** Results from surface plasmon resonance analyses of flagellin interactions with TLR5^N14^. The mean ± standard deviation of the compiled results for each flagellin analysed by SPR. The mean and standard deviation were determined by one-way ANOVA with Šídák correction in Prism 10 (GraphPad).

| Flagellin construct | K<sub>D</sub> [nM] | k<sub>a</sub> [1/Ms] | k<sub>d</sub> [1/s] |
| --- | --- | --- | --- |
| *R. hominis* FlaB | 11.38 ±3.339 | 45157 ±7459 | 5.547e<sup>-4</sup><br>±2.704e<sup>-5</sup> |
| *R. hominis* FlaB-ΔD0 | 7.081 ±0.087 | 75480 ±2424 | 5.344e<sup>-4</sup><br>±1.076e<sup>-5</sup> |
| *R. hominis* FlaB-PIM | 77.99 ±21.15 | 9091 ±1524 | 6.907e<sup>-4</sup><br>±1.289e<sup>-4</sup> |
| *S. typhimurium* FliC | 16.26 ±5.566 | 11293 ±3148 | 1.925e<sup>-4</sup><br>±1.143e<sup>-4</sup> |
| *S. typhimurium* FliC-ΔD0 | 8.772 ±1.012 | 15573 ±357.3 | 1.367e<sup>-4</sup><br>±1.687e<sup>-5</sup> |
| *S. typhimurium* FliC-PIM | 585.1 ±26.08 | 18830 ±1981 | 0.01100<br>±6.834e<sup>-4</sup> |
| *S. typhimurium* FliC-PIM-ΔD0 | 274.3 ±13.76 | 42807 ±1612 | 0.01173<br>±2.740e<sup>-4</sup> |

### *St*FliC and *Rh*FlaB have distinct temporal profiles in TLR5 binding

*St*FliC associated (ka 11293 Ms^-1^) and dissociated (kd 0.0001925 s^-1^) significantly slower than *Rh*FlaB (ka 45157 Ms^-1^, kd 0.0005547 s^-1^; Fig. 2 B,C; Table 1). *St*FliC- ΔD0 showed no significant difference in binding kinetics compared to *St*FliC, while *Rh*FlaB-ΔD0 (ka 75480 Ms^-1^) showed a significant increase in association compared to *Rh*FlaB (ka 45157 Ms^-1^; Table 1; Supp. Fig. 1).

We observed that *Rh*FlaB-PIM (ka 9091 Ms^-1^, kd 0.0006907 s^-1^) associated significantly more slowly than *Rh*FlaB, but showed no significant difference in dissociation rate. *St*FliC-PIM (ka 18830 Ms^-1^, kd 0.01100 s^-1^) both associated and dissociated significantly faster than *St*FliC, though the difference in association was comparatively marginal (Table 1; Fig. 2B-C). These differences in kinetic binding profiles for *Rh*FlaB and *St*FliC are apparent in the SPR binding curves (Supp. Fig. 2). Our results suggest that for *Rh*FlaB, the PIM residues are essential for TLR5 binding. However, the PIM residues are not necessary for association in *St*FliC, and instead appear to stabilise the *St*FliC:TLR5^N14^ complex.

### Cryo-EM structure of the *Rh*FlaB flagellum complex and flagellin monomer

We used cryo-EM to resolve the structure of the *Rh*FlaB flagellin from the native *R. hominis* flagellum, in a manner comparable to an existing cryo-EM derived *St*FliC structure (Yamaguchi et al. 2020). We grew *R. hominis* under anaerobic conditions to stationary phase, then sheared and purified flagella from the cultured cells. We verified that isolated flagellar filaments were composed of *Rh*FlaB using SDS-PAGE and Western blot with an antibody targeting *Rh*FlaB (Ab-*Rh*FlaB) (Supp. Fig. 3). Both analyses showed two distinct bands at ∼60 kDa and ∼40 kDa in fractions 5-8, which differed from the ∼54 kDa band predicted for *Rh*FlaB using the ExPASy Compute pI/Mw tool (Gasteiger et al. 2005). However, as no cross-reactivity was observed for the antibody, all other flagellins identified from the *R. hominis* genome were predicted to be >30 kDa, we concluded that our samples were primarily composed of *Rh*FlaB.

To confirm the purity and distribution of the purified flagella sample, we used negative staining (Supp. Fig. 4A). We recorded 2,480 TEM images of *R. hominis* flagellar filaments, then manually filtered the images for quality (Supp. Fig. 4B).

Through auto-picking, we extracted 576,585 particles, which we sorted by 2D class averaging to a final dataset of 443,605 flagella particles (Supp. Fig. 4C). We identified twist (65.4424°) and rise (4.69Å) parameters, facilitating 3D helical refinement. This calculated a final 3.1Å electron density map of the supercoiled flagella, with low variation in resolution throughout the structure (Fig. 3A; Supp. Fig. 5).

**Figure 3.**
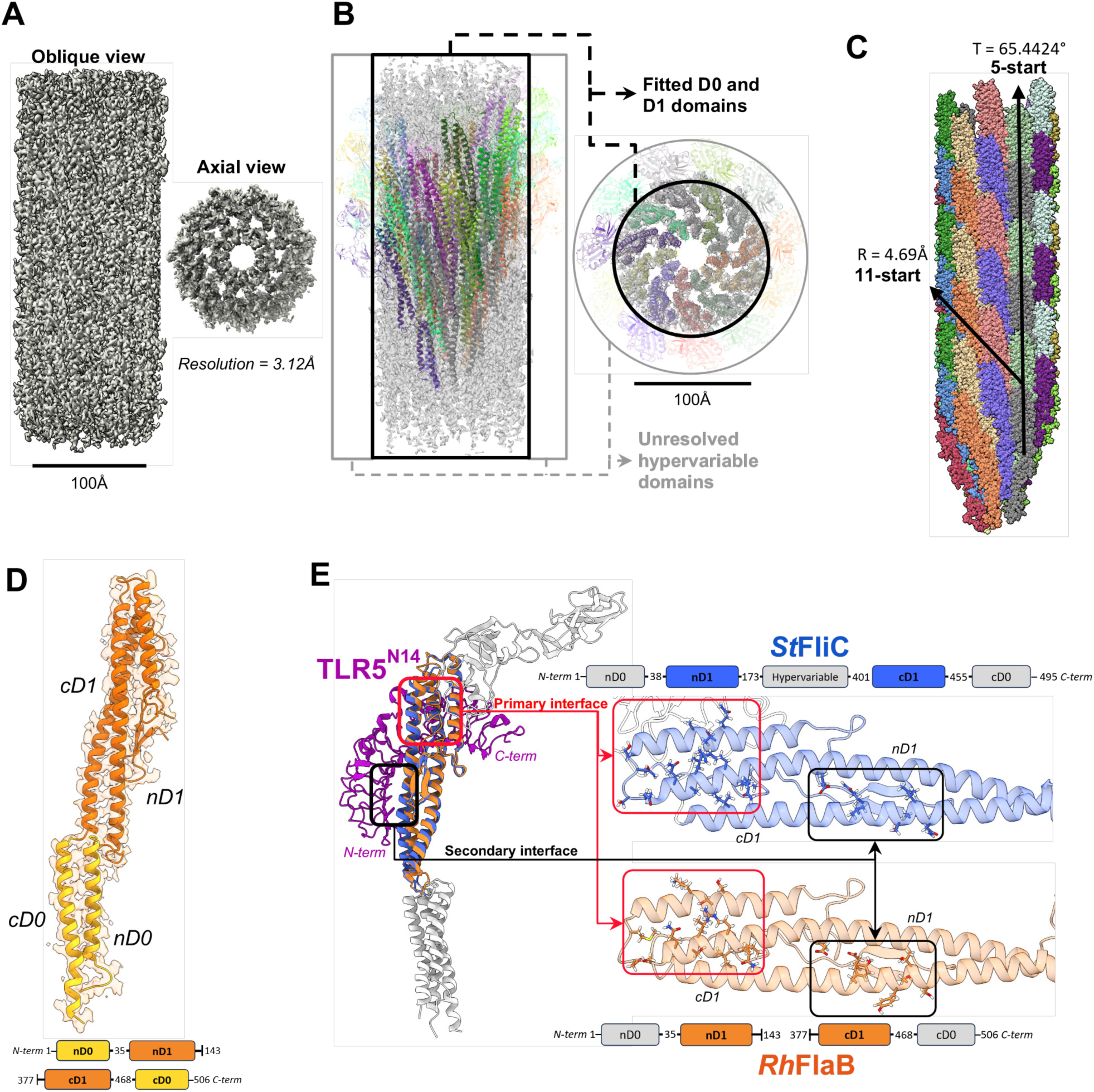
Cryo-EM structure of the *R. hominis* flagella and flagellin monomer. (A) Oblique and axial view of the refined *R. hominis* flagellum density map. The filament segment has a diameter and length of ca. 123 Å and 270 Å respectively. The reported resolution of the structure is 3.12 Å (Supp. Fig. 5). (B) Fitted 22-flagellin *Rh*FlaB complex in the post-processed density map. D0 and D1 domains fit in the density map, but densities corresponding to the hypervariable domains were not resolved. (C) 3D atomic reconstruction of the *Rh*FlaB flagellum segment, without the hypervariable domains. Repeated flagellins are shown in the same colour. The rotational symmetry, twist, and rise, are indicated on the appropriate axis. (D) The D0 and D1 domains of a *Rh*FlaB monomer fitted into the electron density map, coloured according to the domain map. (E) Superposition complex of *Rh*FlaB and *St*FliC (PDB ID #6jy0) bound to TLR5^N14^. The TLR5^N14^ structure was generated with AlphaFold2 (Jumper et al. 2021). The primary and secondary interfaces are indicated. TLR5 binding residues in *St*FliC absent from *Rh*FlaB are shown (blue). The corresponding *Rh*FlaB residues (orange) were identified by sequence and structure alignment (Supp. Fig. 6). An annotated overview of each interface is shown in Supp. Fig. 7.

We modelled *Rh*FlaB monomer structures with AlphaFold2 and fitted them into the density map, building a 22-flagellin complex (Jumper et al. 2021). The fitting showed that the resolved density map covered the D0 and D1 domains, but did not cover the hypervariable domains (Fig. 3B). This absence may stem from the flexible and unstructured nature of these domains. However, the two protein bands in the SDS-PAGE and western blots also indicate that the ∼24 kDa hypervariable domains were degraded in a significant portion of the flagella, which would further average these densities out of the final reconstruction (Supp. Fig. 3). We excluded these domains from the model prior to refinement.

Individual flagellin monomers were refined to solve the map, then built into the final flagellum complex (Fig. 3C). This complex forms an 11-start right-handed helix, common for flagellins displaying the TLR5 epitope, such as *St*FliC (Croll 2018; Mimori et al. 1995; Blum et al. 2019). The final *Rh*FlaB structure showed a good fit to the density map (Fig. 3D), no severe clashes, and no residues outside of energetically allowed regions (PDB ID: ####).

### Residues involved in *St*FliC - TLR5 interactions cluster in two distinct groups in the D1 domain

Previous studies of flagellin-TLR5 interactions have identified several D1 domain binding residues in *St*FliC (Song et al. 2017; Yoon et al. 2012; Smith et al. 2003; Ivičak-Kocjan et al. 2018; Forstnerič et al. 2016). Through sequence and structural alignments, we identified the residues in *Rh*FlaB which differed from *St*FliC D1 domain TLR5 binding residues (Supp. Fig. 6). When mapped onto an existing *St*FliC structure (PDB ID #6jy0 from (Yamaguchi et al. 2020)) and our *Rh*FlaB structure, we observed that these residues clustered into two distinct regions, which we refer to as the primary interface (PI) and secondary interface (SI) (corresponding to Interfaces B and A in (Yoon et al. 2012; Song et al. 2017)) (Fig. 3E; Supp. Fig. 7).

We superimposed the *St*FliC and *Rh*FlaB structures on an AlphaFold2 generated model of TLR5^N14^, to create *in-silico* TLR5^N14^:*St*FliC and TLR5^N14^:*Rh*FlaB complexes. We then aligned the superposition complexes based on an existing crystal structure of zebrafish TLR5b (*dr*TLR5b) in complex with a truncated *Sd*FliC (PDB ID #3v47 from (Yoon et al. 2012)) (Fig. 3E). Based on the buried solvent-accessible surfaces in the superposition complexes, the PI and SI of both flagellins were predicted to interact with TLR5^N14^.

### *Rh*FlaB and *St*FliC show distinct differences in surface hydrophobicity at both D1 binding interfaces

The flagellin PI lies above a binding pocket in TLR5 formed by the leucine rich repeat 9 (LRR-9) loop (Yoon et al. 2012). The PI surface constitutes a single hydrophobic pocket on *St*FliC, whereas it forms two larger pockets on *Rh*FlaB (Fig. 4A,B). The shared hydrophobic pocket is formed by a leucine and isoleucine residue (*St*FliC: L94, I111; *Rh*FlaB: L93, I111). Based on the superposition complexes, we propose that they interact with TLR5: I257. The second hydrophobic pocket in *Rh*FlaB is primarily formed by I86 and L118, and is predicted to interact with TLR5: F282. The two hydrophobic pockets on *Rh*FlaB bracket R89 and Q88, and neighbour Q96 (*St*FliC: R90, Q89, Q97 respectively), which comprise the three TLR5 epitope residues mutated in the PIM flagellins, form hydrogen bonds at the LRR-9 loop binding pocket (Fig. 4C) (Yoon et al. 2012; Song et al. 2017).

**Figure 4.**
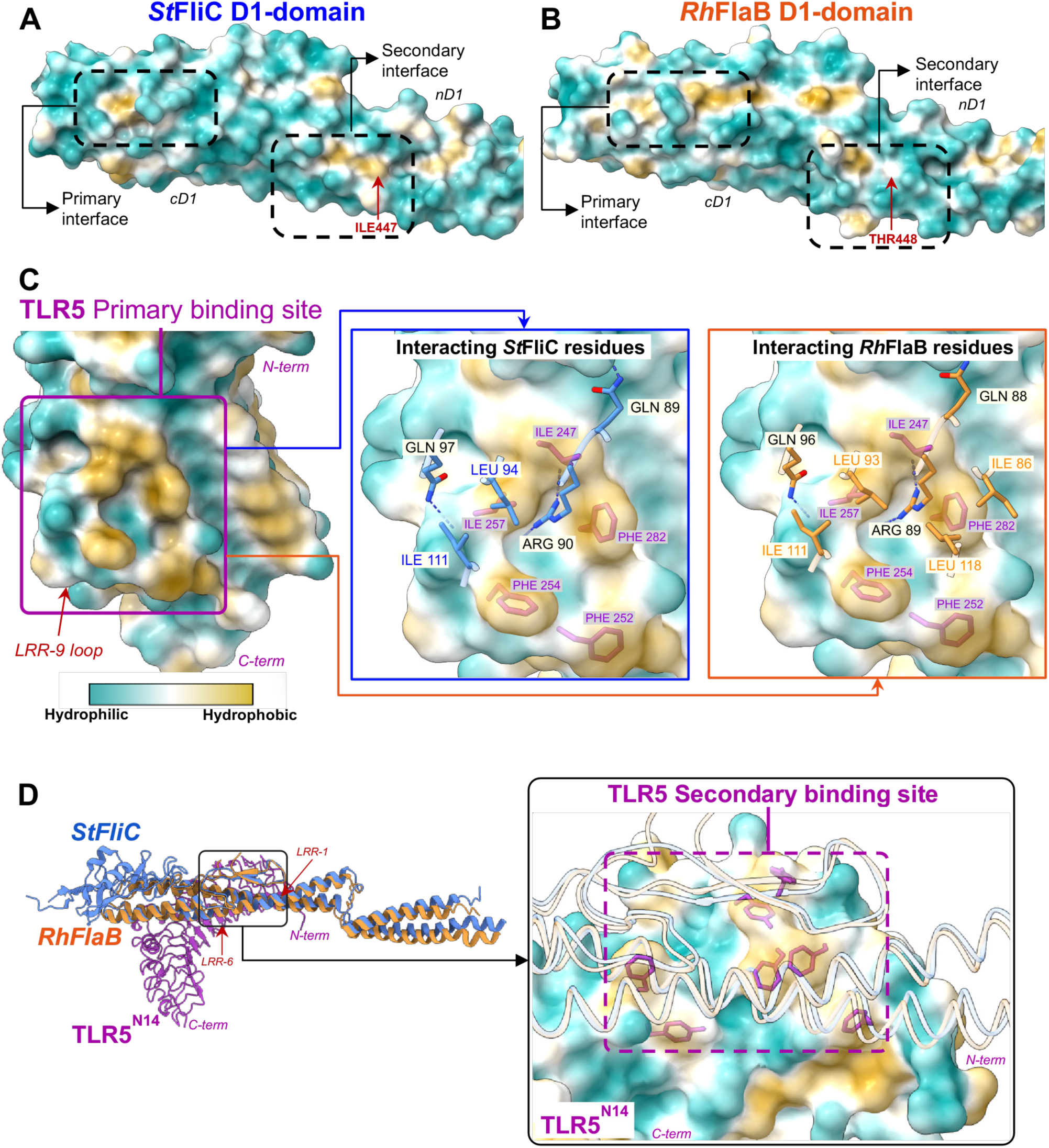
Comparison of hydrophobic surface profiles at the primary and secondary binding interfaces. (A-B) Hydrophobic surface profiles for *St*FliC and *Rh*FlaB D1-domain binding interfaces. The region comprising the PI and SI are outlined. (C) Overview of the TLR5 PI binding site (purple), comprised of the LRR-9 loop and neighbouring residues. The *St*FliC (blue) and *Rh*FlaB (orange) residues expected to interact are displayed, with notable hydrophobic residues labelled in blue/orange, and the hydrogen-bond forming ‘PIM’ residues labelled in black. Hydrophobic TLR5 residues predicted to interface with each flagellin are labelled (purple). (D) The TLR5 SI binding site predicted to interact with flagellin based on the superposition complex. Hydrophobic residues predicted to associate with the flagellins are displayed. *St*FliC (blue) and *Rh*FlaB (orange) structures are overlaid for context.

*St*FliC has two hydrophobic pockets at the SI; the largest is located between the nD1 and cD1 ɑ-helices. A second hydrophobic pocket is located on the converse cD1 ɑ-helix surface and contains an isoleucine (*St*FliC: I447) implicated in TLR5 binding (Fig. 4A) (Smith et al. 2003; Song et al. 2017). While corresponding hydrophobic pockets are present on *Rh*FlaB, they are reduced in scope. The cD1 ɑ- helix notably lacks an isoleucine corresponding to *St*FliC: I447, containing a threonine in its place (*Rh*FlaB: T448) (Fig. 4B). The corresponding binding interface on TLR5^N14^ is distributed between LRR-1 and LRR-6 (Fig. 4D). This region contains numerous aromatic amino acids such as phenylalanines, which are predictive of protein-protein binding sites, but a lack of existing structural data means that direct interactions cannot be predicted at this interface (Bordner 2009; Kotha and Staller 2023).

### *Rh*FlaB displays a highly negatively charged surface across D1 domain interfaces

Electrostatic interactions between individual residues were particularly difficult to analyse with superposition complexes, as direct charged interactions could not be extrapolated. Instead, we explored a general view of the surface charge profiles across the binding interfaces.

*St*FliC displays a mix of negative, neutral, and positively charged residues at the PI. The SI is primarily neutral, with few negatively charged residues (Fig. 5A). Interestingly, *Rh*FlaB shows a distinct strongly negatively charged surface across both TLR5 binding interfaces (Fig. 5B). This was most pronounced at the SI, which displays a near-completely negatively charged surface. The corresponding TLR5 interfaces display primarily neutral surface charges, resulting from the abundance of non-polar phenylalanine residues (Fig. 5C). Consequently, no pockets of charged residues were present on TLR5 that could be expected to favour binding with the negatively charged *Rh*FlaB surface.

**Figure 5.**
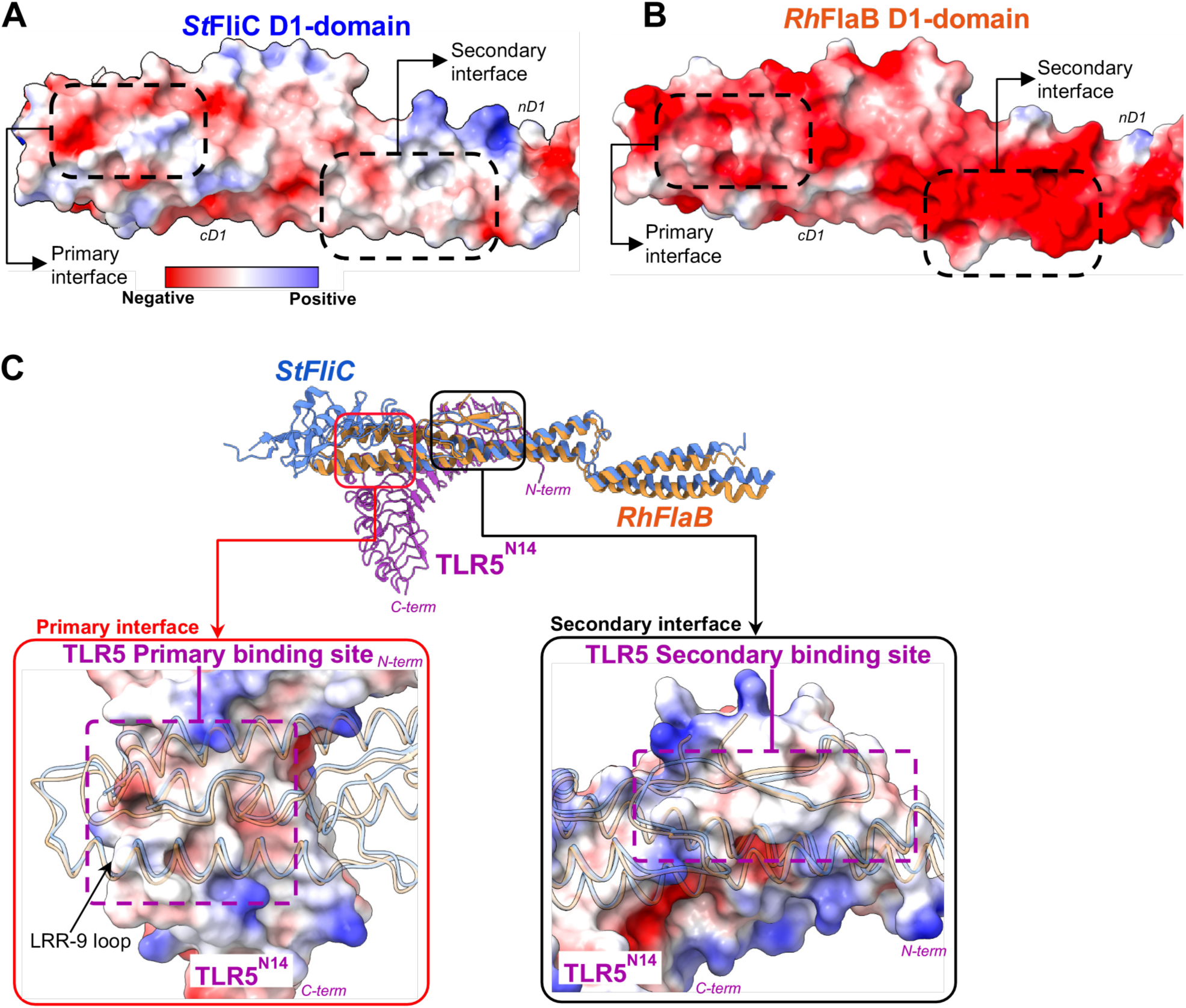
Comparison of electrostatic surface profiles at the primary and secondary binding interfaces. (A-B) Electrostatic surface profiles for *St*FliC and *Rh*FlaB D1-domain binding interfaces. The region comprising the SI and PI are indicated. (C) Superposition complex showing the corresponding TLR5 Primary (red) and Secondary (black) TLR5 binding sites predicted to interface with flagellin. Interfacing regions are comprised of numerous hydrophobic residues, resulting in a primarily neutrally charged surface profile. *St*FliC (blue) and *Rh*FlaB (orange) structures are overlaid for context.

## Discussion

Here, we provide additional insight into the mechanism by which flagellins are silent. We show that while displaying a low KD, the silent flagellin *Rh*FlaB remains bound at its D1 domains to the TLR5 canonical binding site for only a short period compared to the stimulatory flagellin *St*FliC. We propose that this rapid dissociation of the silent flagellin may limit the ability for TLR5 to trigger a downstream immune response.

We had recently described flagellins from commensal human gut bacteria that exhibit a “silent” phenotype in their interaction with TLR5. The D1 domains of silent flagellins bind TLR5 strongly at the canonical binding site, yet the receptor response is muted (Clasen et al. 2023). We showed that silent flagellins lack a previously uncharacterized allosteric binding site on the D0 domain that allows stimulatory flagellins such as *St*FliC to access preformed dimers on the cell surface, and that this allosteric binding site endows *St*FliC with its characteristic strong agnosim (Clasen et al. 2023). However, the D0 domain alone is insufficient to turn a silent flagellin into a stimulatory flagellin, implying that other interactions on the D1 are also in play that remain to be characterized. The combined results shown here indicate that binding kinetics at the D1 may also play a role in the silent flagellin phenotype.

Our SPR-generated KD measurements corroborate our previous observations based on relative binding: *Rh*FlaB binds TLR5 more strongly than *St*FliC, and PIM mutations severely impair binding in both (Clasen et al. 2023). The slower dissociation that we observed in *St*FliC indicated that it bound TLR5^N14^ for a substantially longer duration than *Rh*FlaB, despite the slow association rate. These results suport *St*FliC’s ability to form a comparatively more stable complex with TLR5.

The substantial increase in KD seen in the PIM mutants compared to the WTs stemmed from different aspects of their kinetic binding profile. *Rh*FlaB-PIM had a very low ka compared to *Rh*FlaB, which was expected based on the purported role of the TLR5 epitope residues in complex formation, but showed no significant change in kd ^(^Song et al. 2017; Smith et al. 2003^)^. Conversely, *St*FliC-PIM surprisingly had comparable ka compared to *St*FliC, but instead had a dramatic increase in kd. Taken together, these findings indicate that *Rh*FlaB is reliant on the PI for TLR5^N14^ binding, with no strong interactions occurring at the *Rh*FlaB SI. *St*FliC, however, associates at a region distal to the PI, likely at the SI, and instead utilises the TLR5 epitope as an anchor to stabilise the complex.

The two hydrophobic pockets found on the *Rh*FlaB PI surface, which bracket the three hydrogen-bond forming ’PIM’ residues, likely improves the ability of *Rh*FlaB to associate in an orientation which encourages hydrogen-bond formation between the TLR5 epitope and the TLR5 LRR-9 loop binding pocket (Song et al. 2017; Yoon et al. 2012). Conversely, the single hydrophobic site in the *St*FliC PI decreases the likelihood that the flagellin will bind in such an orientation, resulting in the lower ka. Thus, the kinetics of flagellin association are likely determined by the conformational changes of the binding residues at the TLR5 epitope.

While an existing structure has explored interactions at the SI in *Sd*FliC, it displays artefacts from the use of a truncated flagellin, which prevented us from investigating direct interactions between residues at this interface (Yoon et al. 2012). Instead, we derived results from more general characteristics of the SI. The *St*FliC SI contains two broad hydrophobic pockets, which align over hydrophobic phenylalanine residues on the corresponding TLR5 binding site. While these were also present in *Rh*FlaB, they were smaller in scope and occluded by protruding hydrophobic residues. We predict that hydrophobic aromatic residues on the TLR5 surface more readily interact with hydrophobic pockets in *St*FliC, evidenced by the presence of *St*FliC: I447, a known binding residue, in one of these pockets (Smith et al. 2003). However, hydrophobicity alone did not clearly explain the apparent lack of binding at the *Rh*FlaB SI.

While both flagellins share a primarily negatively-charged PI, the highly negatively charged *Rh*FlaB SI is a dramatic contrast. In the flagellum, these negatively charged surfaces are oriented towards the hollow centre, and may play a role in protein trafficking as in other helical complexes (Gatsogiannis et al. 2016; Feld, Brown, and Krantz 2012). However, this charged surface conveys no clear advantage when binding to TLR5, and may instead negatively impact diffusion and binding potential (Xu et al. 2013; Estrada et al. 2023; W. He et al. 2020; Farías-Rico et al. 2018).

We propose that the negatively charged *Rh*FlaB SI limits its ability to bind TLR5 compared to the neutrally charged *St*FliC. This would explain the inability of *Rh*FlaB-PIM to bind TLR5^N14^, and the comparatively lower stability of the TLR5^N14^:*Rh*FlaB complex. These results imply that the ability to bind at both D1 domain interfaces, not just the PI, is necessary for the formation of a stable complex capable of promoting a TLR5 immune response.

The limitations of our study include that we generalize properties of stimulatory and silent flagellins from one example of each, with the caveat that other flagellins may interact differently with the TLR5 receptor. Our analyses were conducted in vitro, which may differ from in-cell conditions, and with a truncated version of the TLR5 ectodomain, which may differ from the full-length. Further studies in which a wider range of flagellins are tested on full-length TLR5 ectodomain will allow us to assess the generalities of the findings presented here.

Taken together, our findings shed light on how binding at the D1 domain can facilitate silent, stimulatory, and evasive responses when binding TLR5. These outcomes are primarily determined by binding kinetics, and not KD. In light of our previous findings, our results show how the D1 and D0 domains may work in concert to modulate TLR5 activity, and build the foundation for further kinetic studies investigating flagellin D0 domain interactions (Clasen et al. 2023). The identification of structural motifs driving the kinetic binding profiles can inform the study of further bacterial flagellins, and be applied in the design of ligands targeting TLR5.

## Methods

### Recombinant flagellins

Wild-type flagellin sequences for StFliC, and RhFlaB were obtained from Uniprot (ID #P06179, and #G2T5W2 respectively). PIM mutants (StFliC: Q89A, R90A, Q97A; RhFlaB: Q88A, R89A, Q96A) previously observed to abrogate binding (Song et al. 2017; Yoon et al. 2012; Smith et al. 2003; Ivičak-Kocjan et al. 2018; Forstnerič et al. 2016), were included, as previously described (Clasen et al. 2023), as well as D0-truncated flagellins. Codon-optimised plasmids containing the recombinant flagellins were produced in *E. coli* as previously described (Clasen et al. 2023). For purification, cell pellets were resuspended in 1:10 w/v of resuspension buffer (20 mM Tris-HCl, 300 mM NaCl, 5 mM Imidazole, 1% TritonX-100, pH 7.5, protease inhibitor (cOmplete, EDTA-free, Roche #5056489001)), then sonicated at least 5 times with a SONOPULS mini20 ultrasonic homogenizer (BANDELIN) (1 min, 5 s pulses, 40% amplitude). Lysed cells were cleared by centrifugation (30 min, 18,000 rcf, 4°C) and filtered. Columns containing nickel-nitrilotriacetic acid (Ni-NTA) agarose (QIAGEN #30250) were equilibrated with 2 column volumes (CV) of wash buffer (20 mM Tris-HCl, 300 mM NaCl, 20 mM Imidazole, pH 7.5). Cleared lysates were incubated with Ni-NTA agarose in the column for 1 hour at 4 °C, then flowed through. The resin was washed with 2 CV wash buffer, then eluted with 0.5 CV elution buffer (20 mM Tris-HCl, 300 mM NaCl, 300 mM Imidazole, pH 7.5).

The eluent was concentrated to 1 mL with Pierce 10K MWCO protein concentrators (Thermo Fisher Scientific #8852) and further purified by size exclusion chromatography (SEC) with an Äkta Pure protein purification system (Cytiva) equipped with a Superdex® 200 Increase 10/300 GL column (Cytiva) in SEC buffer (300 mM NaCl, 50 mM HEPES, 1 mM TCEP, 5% Glycerol, pH 7.5). The eluent was collected in a 96-well plate, and verified by SDS-PAGE. Protein samples were then pooled, and the concentration measured (DS-11+ Spectrophotometer, DeNovix). All samples were stored at -20°C.

### Purification of TLR5^N14-^

The production of TLR5^N14^ was performed following the protocol described in (Clasen et al. 2023). In brief, IgG-FC-tagged TLR5^N14^ was synthesised with an N-terminal GP67 secretion sequence (Kretzschmar et al. 1996), and cloned into a modified pLIB vector containing a C-terminal 6xHis affinity tag. The vector was cloned into DH10EMBaCY cells to generate a recombinant baculovirus genome (bacmid). SF9 insect cells were inoculated with the bacmid, and the virus expanded through two rounds of amplification to a ‘V2’ expression culture. Protein production was performed in Hi5 cells using a 1:100 (v/v) inoculation of V2 supernatant. After 72 hours, the supernatant was harvested by centrifugation (800 rcf, 30 minutes, 4°C) and filter purified (Steritop 0.22 µm 250 mL Threaded Bottle Top Filter; Millipore # SCGPS02RE).

The supernatant was affinity purified using an Äkta Start protein purification system (Cytiva) equipped with two HisTrap HP 5 mL protein purification columns (Cytiva #175248). The supernatant was run through the HisTrap columns, which were then washed with 12 CV of wash buffer. Samples were eluted with 8 CV elution buffer in 4 mL fractions, and expression checked by SDS-PAGE. A final SEC purification step was performed with the same methodology as for flagellin purification. Samples were pooled and stored at -20°C.

### Surface Plasmon Resonance

Flagellin samples were dialyzed in 1L of PBS- T (pH 7.5) at 4°C twice for ≥1 hour, and a third time overnight, in Slide-A-Lyzer 10K MWCO dialysis cassettes (Thermo Fisher Scientific #66380). Following the third dialysis, 500 mL of dialysis buffer was collected, filtered, and degassed for use as the SPR running buffer. Each SPR running buffer was used only for the samples it was used to dialyse. The protein concentration was measured, the sample diluted to the highest concentration (25 ng/μL for RhFlaB constructs, 50 ng/μL for StFliC constructs) in SPR running buffer, then a fivefold 1:2 dilution series was made for single-cycle analysis.

SPR was performed using a Biacore X100 (Cytiva) equipped with a Sensor Chip Protein A (Cytiva #29127558). Pure TLR5^N14^ was defrosted on ice, diluted to 175 ng/μL in SPR running buffer, then captured for 800 seconds. Run conditions were optimised for *St*FliC (360 s contact, 240 s dissociation, 15 μL/min flow rate) and *Rh*FlaB (180 seconds contact, 240 seconds dissociation, 30 μL/min flow rate), with mutants run at the same conditions as their respective WT. Only two samples could be measured per run, so each run paired WT *St*FliC or *Rh*FlaB against each-other or a respective mutant. SPR running buffer measured as a blank reference under the same run conditions, and subtracted from the sample curve. Three technical replicates were performed per sample per run.

Curve fitting and analysis were performed with the Biacore X100 Evaluation Software (Cytiva). The affinity constant (ka), dissociation constant (kd), and binding strength (KD) were determined for each replicate. KD values shown were calculated based on the reaction rate constants ka and kd, due to the relevance of the kinetics constants in this study. All curves passed the software’s quality-control metrics, and the closeness of the fitted curve was determined by the Chi square and Rmax values. Individual curves displaying poor immobilisation of TLR5^N14^ were omitted from analysis.

### Statistical analysis

Statistical analysis was performed with Prism 10 (GraphPad). Kinetics data are presented as arithmetic means ± standard deviations (SD). Statistical significance was determined with a one-way analysis of variance (ANOVA) with Šídák correction in Prism 10 (GraphPad). P-values < 0.05 were considered statistically significant.

## Cryo-EM of *R. hominis* flagellar filaments

### R. hominis cell culturing

R. hominis cell culturing was performed under anaerobic conditions. R. hominis (DSM #16839) pellets stored in liquid nitrogen were defrosted on ice, then transferred to 10 mL BHIS (DSM #215c) at room temperature. The sample was grown for 24 hours at 37°C, and split to inoculate four Balch tubes containing 10 mL YCFA medium (modified) (DSM #1611) at 1:1000 (v/v) concentration. The samples were cultured at 37°C for 16 hours.

### R. hominis flagella shearing and purification

Shearing was performed in aerobic conditions. One tube (10 mL) of culture was transferred to a 50 mL tube, passaged 15 times through a 18G syringe (Agani #123-3035309), then transferred to a 15 mL tube. This was repeated for the remaining three cultures. Cells were cleared through two rounds of table-top centrifugation (2,800 rcf, 30 min, 4°C), with the supernatant transferred to a clean tube between runs. The supernatant was transferred to 4.7 mL OptiSeal Polypropylene Tubes (Beckman Coulter #361621). The remaining material was pelleted by ultracentrifugation using a TLA-110 rotor and Optima MAX-XP Ultracentrifuge (Beckman Coulter) (100,000 rcf, 1 hr, 4°C). The supernatant was discarded, and the pellet resuspended in 200 μL PBS. Gradient centrifugation was performed in 5 mL Open-Top Thinwall Polypropylene Tubes (Beckman Coulter # 326819). A continuous 20%-60% sucrose gradient in PBS was made using a Gradient Master 108 (Biocomp), and cooled to 4°C. The resuspended pellet was carefully added atop the gradient, then ultracentrifuged using a SW55-Ti rotor and Optima L-100 XP Ultracentrifuge (32,400 rpm (Approximately 127,000 rcf), 3 hr, 4°C). 500 μL fractions were manually aliquoted by pipetting, then checked by SDS-PAGE.

A serum containing a *Rh*FlaB-targeting antibody (Ab-*Rh*FlaB) was generated, and checked for cross-reactivity against other recombinant flagellins from the *R. hominis* genome, *St*FliC, and *E. coli* cell lysate. Minimal cross-reactivity was observed (Supp. Fig. 8). Western blotting was used to confirm that fractions containing protein bands seen by SDS-PAGE corresponded to *Rh*FlaB. Fractions with the highest protein concentration were pooled and transferred to 4.7 mL OptiSeal Polypropylene Tubes, and ultracentrifuged (TLA110 rotor, 150k rcf, 1 hr, 4°C). The sample pellet was washed in PBS, pelleted again, then resuspended a final time in 100 μL PBS. The concentration of each sample was determined with the Pierce BCA Protein Assay Kit (Thermo Fisher Scientific #23225).

### Negative staining

Flagella samples were imaged after negative staining (NS) for quality control prior to cryo-EM. Samples were diluted to a concentration of 50 ng/μL with PBS and negatively stained with 1% uranyl acetate on glow-discharged carbon-coated 300 mesh copper grids. Samples were imaged by TEM using a Tecnai Spirit (Thermo Fisher Scientific). Representative images showing flagellin concentration and distribution were taken for each sample (Supp. Fig. 4A).

### Cryo-EM imaging and data processing

Purified flagella samples were applied on glow-discharged R1.2/1.3 300 mesh copper grids (Quantifoil) and plunge frozen in liquid ethane (EM GP, Leica). Grids were imaged with a Talos Arctica (Thermo Fischer Scientific) at 200 kV equipped with a Falcon III camera (Thermo Fischer Scientific) operated in linear mode. Data was collected using EPU acquisition software (Thermo Fisher Scientific) at a nominal magnification of 150,000, which corresponds to a calibrated pixel size of 0.97 Å. MotionCorr2 and CTF-correction were performed with TranSPHIRE (Stabrin et al. 2020), then automated filament picking was performed with crYOLO (Wagner et al. 2019).

Extracted particles were imported into Relion (Scheres 2012). An initial model was generated based on a hollow cylinder, and used for 3D refinement with helical reconstruction (S. He and Scheres 2017). Exact twist and rise parameters were determined with local searches of symmetry. Iterative refinements were performed using 3D refinement maps as the initial model. Post processing, CTF refinement and Bayesian polishing were performed to calculate the final density map.

### Structure fitting & refinement

ChimeraX and ISOLDE were used for fitting, model building, and visualisation (Croll 2018; Pettersen et al. 2021). An existing 22- flagella complex structure (PDB ID #6jy0 from (Yamaguchi et al. 2020)) was fitted into the density map. An initial RhFlaB structure was generated with AlphaFold2 based on the type strain (DSM16839) (Jumper et al. 2021; Travis et al. 2015; MacDougall et al. 2020). RhFlaB monomers were fitted based on the fitted 6jy0 complex, producing a 22-flagellin RhFlaB complex. A central flagellin with the most inter-protein interactions was refined, then used to rebuild the complex. This process was repeated until no further clashes appeared when rebuilt. The structure was then refined as a monomer to resolve remaining outliers, for use in the superposition complex.

## Comparison and analysis of protein structures

### Identification of TLR5-binding regions of StFliC and RhFlaB

We searched for StFliC residues experimentally shown to significantly impact the human TLR5 innate immune response (Song et al. 2017; Yoon et al. 2012; Andersen-Nissen et al. 2007; Jacchieri, Torquato, and Brentani 2003; Murthy et al. 2004) in RhFlaB by comparins AA sequences using Clustal O (1.2.4) (Supp. Fig. 6). We identified 21 residues that differed between StFliC and RhFlaB, and mapped them onto the StFliC and RhFlaB structures to verify their structural alignment (Fig. 3E; Supp. Fig. 7).

### Analysis of binding interfaces

Superposition complexes were made using ChimeraX, to examine how the experimental structures and binding interfaces aligned with TLR5^N14^. An existing comparable complex, Zebrafish TLR5b (zTLR5b):StFliC (PDB ID #3v47 from (Yoon et al. 2012)), was used for reference. A TLR5^N14^ structure generated with AlphaFold2 was fitted to zTLR5b. The RhFlaB monomer structure and a comparable experimental StFliC monomer (PDB ID #6jy0 from (Yamaguchi et al. 2020)) were fitted based on the corresponding flagellin in the zTLR5b:StFliC complex. The superposition showed that the alignment of the flagellins was imperfect, and the flagellin from the complex appeared to differ in confirmation. Consequently, the experimental RhFlaB and StFliC structures were both fitted either based on the alignment of theTLR5 epitope, or the secondary interface, as appropriate for the analysis.

The binding interfaces were analysed in the context of their alignment in superposition complexes. The corresponding interfaces on TLR5^N14^ were identified based on predictions from ChimeraX. For the Primary interface, the PIM residues were adjusted using ChimeraX, to reflect the hydrogen-bond forming rotamer orientations seen in zTLR5b (Yoon et al. 2012). H-bonds were simulated to confirm the same interactions were expected in human TLR5 (Fig. 4C). Hydrophobic and electrostatic surface profiles were generated using ChimeraX to analyse the binding interfaces.

## Acknowledgments

We thank Andrei Lupas and members of the Ley Lab for their assistance. This work was funded by the Max Planck Society. The authors declare no competing interests.

